# Genetic tuning of intrinsically photosensitive retinal ganglion cell subtype identity to drive visual behavior

**DOI:** 10.1101/2024.04.25.590656

**Authors:** Marcos L. Aranda, Jacob D. Bhoi, Omar A. Payán Parra, Seul Ki Lee, Tomoko Yamada, Yue Yang, Tiffany M. Schmidt

## Abstract

The melanopsin-expressing, intrinsically photosensitive retinal ganglion cells (ipRGCs) comprise a subset of the ∼40 retinal ganglion cell types in the mouse retina and drive a diverse array of light-evoked behaviors from circadian photoentrainment to pupil constriction to contrast sensitivity for visual perception. Central to the ability of ipRGCs to control this diverse array of behaviors is the distinct complement of morphophysiological features and gene expression patterns found in the M1-M6 ipRGC subtypes. However, the genetic regulatory programs that give rise to subtypes of ipRGCs are unknown. Here, we identify the transcription factor Brn3b (Pou4f2) as a key genetic regulator that shapes the unique functions of ipRGC subtypes and their diverse downstream visual behaviors.

---

Light information from the retina is relayed by more than 40 types of retinal ganglion cells (RGCs) to more than 40 brain regions to mediate conscious visual perception and subconscious, non-image forming functions (*1–5*). Each RGC type exhibits unique morphophysiological properties, projections, and roles in behavior (*6–8*). However, the cellular mechanisms that give rise to this stunning diversity of RGC structure and function are unknown. The melanopsin (Opn4)-expressing, intrinsically photosensitive retinal ganglion cells (ipRGCs) are a class of RGC that play a key role driving both subconscious, non-image forming behaviors such as circadian photoentrainment and the pupillary light reflex as well as contrast sensitivity for conscious visual perception (*9–14*). Different ipRGC subtypes (M1-M6) selectively drive these behaviors, with each subtype characterized by a distinct complement of morphophysiological properties and melanopsin expression levels (*14*). Together, these subtype-defining features form a representative subsample of the broad array of morphophysiological features, projections, and roles in behavior that combine to define all RGC types, making ipRGCs a compelling system for elucidating the mechanisms that drive RGC diversity (*3, 8*). In this paper, we identify a transcriptional program that finely tunes gene expression to produce the range of ipRGC subtypes necessary for diverse visual behaviors.

## Brn3b regulates ipRGC transcriptional identity

Brn3b (Pou4f2) is a transcription factor important for RGC specification in early embryonic development (*15*). Notably, Brn3b expression is present in newly postmitotic ipRGCs and persists into adulthood, suggesting it may play yet unidentified roles in ipRGC development and function (*16*). We therefore assessed how removal of Brn3b from ipRGCs impacted gene regulation in these cells. Since ipRGCs constitute only ∼0.01% of all retinal cells, we took advantage of the translating affinity purification (TRAP) approach to characterize ipRGC gene expression (*17*). We crossed transgenic mice expressing *Opn4^Cre^*, which selectively expresses Cre-recombinase in ipRGCs, with mice conditionally expressing *Rpl22^HA^* in a Cre-dependent manner (*18*). Purification and sequencing of mRNA bound to HA-Rpl22 labeled ribosomes revealed 2326 transcripts enriched in ipRGCs compared to in the total retina (fig. S1). We next generated a Brn3b conditional knockout (Brn3bcKO) animal (*Opn4^Cre^; Brn3b^cKOAP^*) to eliminate Brn3b from postmitotic ipRGCs. Developmentally, mRNA for Brn3b and the mature RGC marker RBPMS are detectable at embryonic day (E) 12.5 in retinal sections, around the time that ipRGCs exit the cell cycle and become postmitotic (fig. S2) (*19, 20*). Importantly, Opn4 mRNA expression is not detectable until E15.5, indicating that Opn4-dependent excision of the Brn3b gene occurs well after postmitotic ipRGC specification (fig. S2). Using mRNA fluorescent in situ hybridization (FISH), we confirmed that the Brn3b expression found in control retinas is absent in ipRGCs of adult Brn3bcKO retinas (fig. S3). We then crossed the Brn3bcKO mouse line with the conditional *Rpl22^HA^* line and performed TRAPseq using retinas from Brn3bcKO (*Opn4^Cre^; Brn3b^cKOAP^; Rpl22^HA^*) and control (*Opn4^Cre^; Rpl22^HA^*) littermates. We identified 1360 transcripts that were differentially expressed in ipRGCs upon knockout of Brn3b (Fig. 1A), with similar numbers of upregulated and downregulated genes, suggesting that Brn3b regulates the expression of a large group of genes in ipRGCs. By examining previous scRNAseq analyses of available datasets, we found that differentially expressed genes identified in Brn3bcKO retinas often also showed graded expression across ipRGC subtypes (Fig. 1, B and C) (*3*), raising the possibility that Brn3b may be linked to ipRGC subtype identity. Indeed, when we analyzed expression of Brn3b itself in the best-characterized ipRGC subtypes (M1, M2, and M4), we identified graded expression of Brn3b, with M1 ipRGCs expressing the least Brn3b, followed by increasing levels in M2 and then M4 cells (Fig. 1D).

**Fig. 1.**
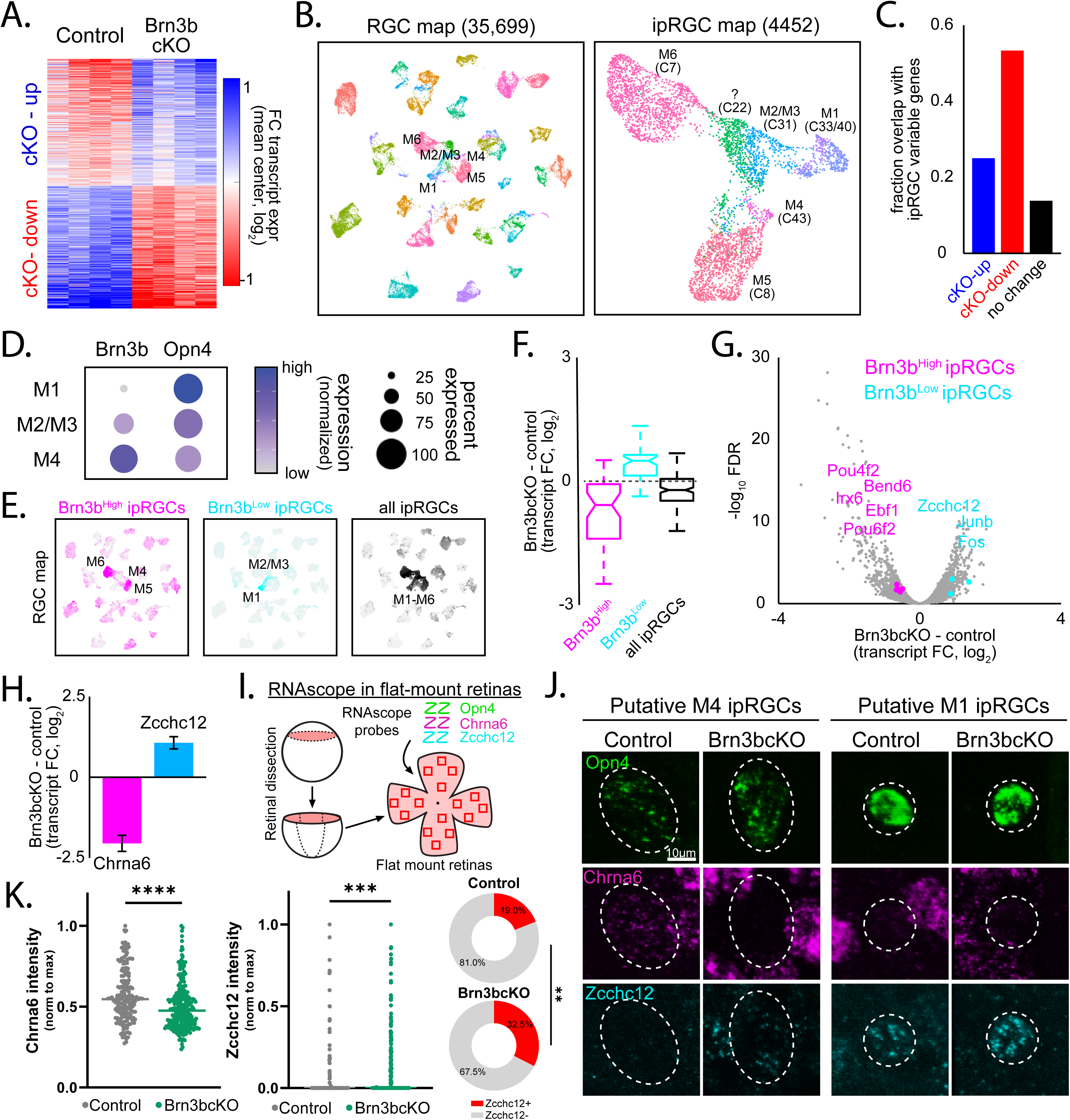
Brn3b confers molecular identity to ipRGC subtypes. (**A**) Differentially expressed transcripts in control and Brn3bcKO ipRGCs. (**B**) ipRGCs identified from publicly available scRNA-Seq profiles of RGCs (*3*), re-analyzed using dimensionality reduction, and visualized with UMAP. Established genetic markers are used to annotate ipRGC subtypes. (**C**) Genes with increased and decreased expression in Brn3bcKO mice exhibited high variability in expression across ipRGC subtypes. (**D**) Brn3b and Opn4 mRNA levels in ipRGC subtypes. Relative expression of Opn4 and Brn3b in M1, M2, and M4 ipRGCs in scRNA-Seq dataset (*3*). (**E**) Top genes that were enriched in M4, M5, and M6 ipRGCs with high levels of Brn3b expression (Brn3b^High^) or in M1 and M2 ipRGC subtypes with low levels of Brn3b expression (Brn3b^Low^) (*3*). (**F**) Brn3bcKO induces opposite effects on transcripts found in Brn3b^low^ and Brn3b^high^ ipRGCs (**G**) Volcano plot of Brn3bcKO-induced changes in transcript expression using TRAP. Transcriptional regulators which define ipRGC subtypes and are dysregulated in Brn3bcKO are highlighted. (**H**) Brn3bcKO-induced changes in Chrna6 and Zcchc12 transcript expression using TRAP. (**I**) Schematic representation RNAscope in flat-mount retinas. (**J**) Representative RNAscope pictures in flat mount retinas showing Chrna6 and Zcchc12 mRNA expression in putative M4 and M1 ipRGCs (dashed ellipses) in control and Brn3bcKO mice. (**K**) Relative expression of Chrna6 and Zcchc12 mRNA in ipRGCs from control and Brn3bcKO mice measured by RNAscope. Lines are median values, **P<0.01, ***P<0.001, Mann Whitney U and Fisher’s exact tests.

Given this differential expression of Brn3b, we next assessed whether Brn3b regulates the transcriptional programs that define ipRGC subtypes. We curated the top genes that were enriched in ipRGC subtypes with high levels of Brn3b expression (M4, M5, and M6 cells, Brn3b^High^) or in ipRGC subtypes with low levels of Brn3b expression (M1 and M2 cells, Brn3b^Low^) using scRNAseq analyses (Fig. 1E). Strikingly, we found that Brn3bcKO retinas showed decreased expression of genes enriched Brn3b^High^ ipRGCs, suggesting that Brn3b controls the transcriptional identity of M4, M5, and M6 ipRGCs. Additionally, we found increased expression of genes enriched in Brn3b^Low^ ipRGCs, indicating that Brn3b negatively regulates the transcriptional identity of M1 and M2 ipRGCs (Fig. 1, E and F). Because groups of transcription factors often work in concert to orchestrate cell identity (*21–23*), we also examined the full set of 11 transcription factors that exhibited graded expression across subtypes and marked Brn3b^High^ or Brn3b^Low^ ipRGCs. Of these, the expression of 8 were dysregulated upon Brn3bcKO (Fig. 1G), suggesting that Brn3b plays a key role in the transcriptional hierarchy that defines ipRGC subtypes.

We further investigated gene regulation by Brn3b across individual ipRGCs using RNAscope. Chrna6, a Brn3b target gene encoding a subunit of the nicotinic acetylcholine receptor, is highly expressed in M4, M5, and M6 ipRGCs but shows only modest expression in subsets of M1 and M2/M3 ipRGCs in control retinas (fig. S1D). Knockout of Brn3b led to reduced Chrna6 expression across all ipRGCs (Fig. 1, H to K), consistent with our observations using TRAPseq. Conversely, Zcchc12, another Brn3b-regulated gene encoding a transcriptional activator, is expressed primarily in M1 ipRGCs in control retinas (Fig. 1G and fig S1D). However, following knockout of Brn3b, we observed an increase in the number of ipRGCs expressing Zcchc12, including putative M4 ipRGCs (Fig. 1, H to K). These findings suggest that Brn3b not only promotes the expression of certain target genes, but it also restricts the expression of other genes to specific ipRGC subtypes, such as M1 cells.

## Brn3b modulates Opn4 expression

Among the genes expressed in ipRGCs, the melanopsin-encoding gene Opn4 is crucial for conferring intrinsic photosensitivity (*9–12, 24*). Moreover, Opn4 expression varies across ipRGC subtypes, with M1 ipRGCs having the highest melanopsin levels, followed by M2, and then M4 cells, in a gradient opposite to that of Brn3b (Fig. 1D) (*13, 24, 25*). The inverse relationship between Brn3b and Opn4 expression across M1, M2, and M4 ipRGCs prompted the hypothesis that the transcription factor Brn3b regulates Opn4’s graded expression patterns across ipRGC subtypes. To test this idea, we quantified Opn4 expression across individual ipRGCs in both retinal sections and retinal whole mounts of adult Brn3bcKO and control littermates using mRNA FISH. We observed a significant increase in Opn4 mRNA expression in Brn3bcKO retinal sections and flat mounts (Fig. 2, A to C and fig S4). Immunohistochemical labeling for Opn4 protein in Brn3bcKO retinas likewise showed higher Opn4 expression, with increased proportions of ipRGCs expressing high levels of Opn4 (Fig. 2, D to F). The total number of ipRGCs remained unchanged (fig. S5). These results indicate that Brn3b plays an important role in suppressing Opn4 expression levels in ipRGCs.

**Fig. 2.**
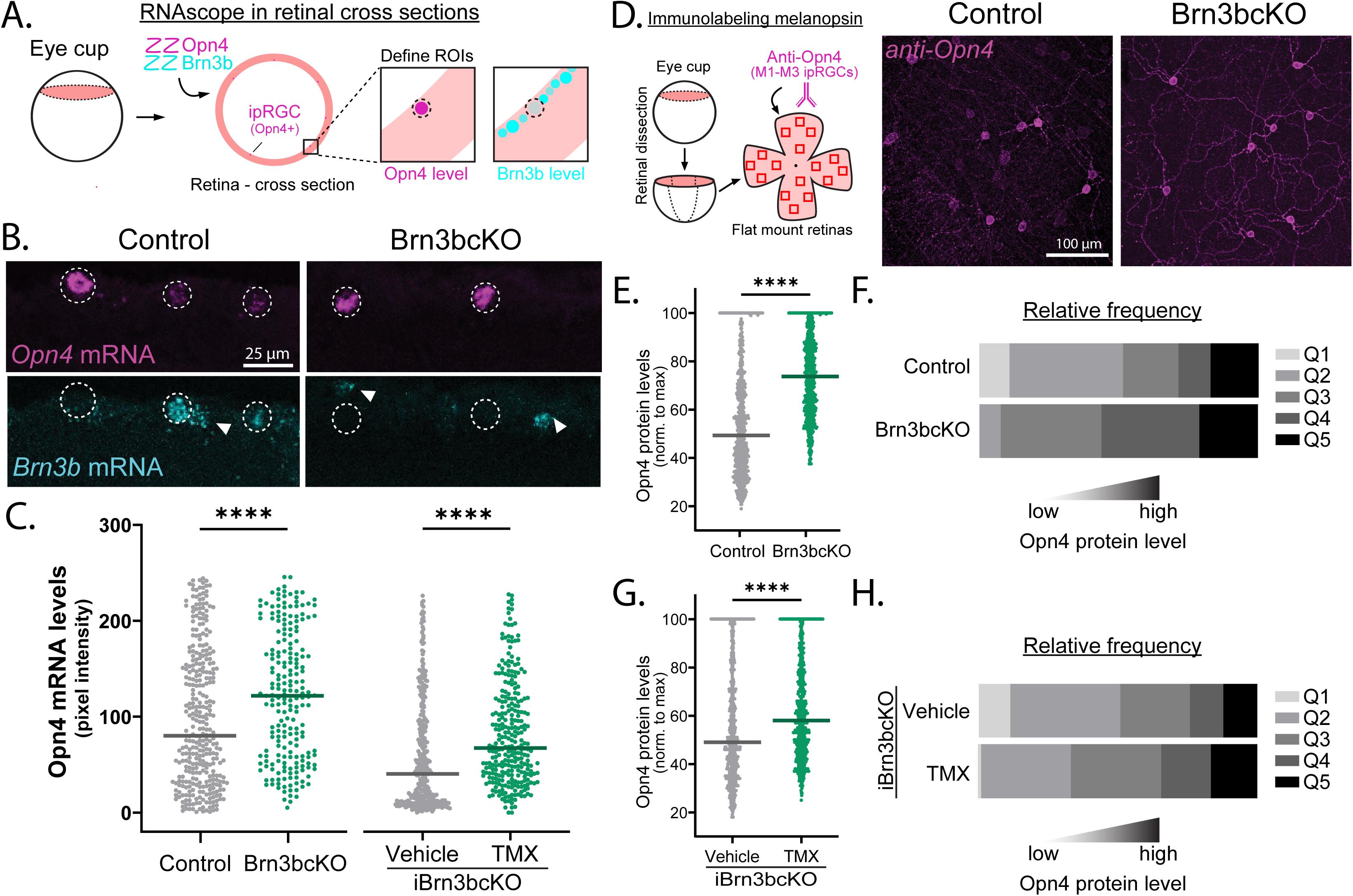
Brn3b represses melanopsin expression. (**A**) Schematic representation of RNAscope retinal cross sections protocol. (**B**) Opn4 and Brn3b mRNA expression in control and Brn3bcKO retinal sections. Brn3b mRNA expression is absent in Brn3bcKO ipRGCs (dashed ellipse) and still present in other RGCs (arrowheads). (**C**) (Left) Opn4 mRNA levels are significantly increased in ipRGCs Brn3bcKO compared to control (n = 216-336 cells/group) littermates. (Right) Opn4 mRNA levels are increased in iBrn3bcKO compared to control ipRGCs (n = 288-371 cells/group). (**D**) (Left) Schematic representation of Opn4 immunolabeling in flat-mount retinas. (Right) Opn4 immunolabeling in ipRGCs of adult control and Brn3bcKO retinas. (**E**-**F**) Opn4 protein levels are increased in Brn3bcKO ipRGCs (E), with an increased proportion of ipRGCs expressing high levels of Opn4 protein (F) (n= 600 cells/group). Q1-Q5: quintiles showing low (Q1) to high (Q5) Opn4 protein levels. (**G**-**H**) Opn4 protein levels are increased in iBrn3bcKO compared to control ipRGCs (G), with an increased proportion of ipRGCs expressing high levels of Opn4 protein (H) (n= 761-800 cells/group). Lines are median values, ****P<0.001, Mann Whitney U test.

In addition to its developmental functions in shaping ipRGC features, Brn3b continues to be expressed in the adult retina where it may continue to affect gene expression (*16*). We therefore tested whether Brn3b regulates Opn4 expression in ipRGCs beyond development by generating a tamoxifen (TMX)-inducible (i)Brn3bcKO line (*Opn4^CreERT2^; Brn3b^cKOAP^*) to remove Brn3b from ipRGCs during adulthood. TMX injection over five consecutive days into adult P60 iBrn3bcKO animals resulted in a significant reduction at P120 of Brn3b mRNA in ipRGCs compared to animals treated with vehicle control (fig. S6). Strikingly, Opn4 mRNA levels were significantly increased in ipRGCs of P120 iBrn3bcKO animals compared to those of control littermates (Fig. 2C). Immunohistochemical labeling also showed an increased proportion of ipRGCs expressing high levels of Opn4 protein upon TMX-induced knockout of Brn3b in adult animals (Fig. 2, G and H). These data suggest that Brn3b is required for maintaining ipRGC subtype-specific Opn4 expression in the adult retina. Collectively, these findings reveal that Brn3b fine-tunes the gene expression profiles of ipRGC subtypes to shape their identity during development and through adulthood.

## Brn3b is a central regulator of morphological features that define ipRGC subtypes

Cellular morphology is a key distinguishing feature of ipRGC subtypes, which we observed to also correlate with Brn3b expression levels. M4 ipRGCs express high levels of Brn3b and have large somata and large, complex dendritic arbors (Fig. 1D, (*12, 26*)). M1 ipRGCs, on the other hand, express low levels of Brn3b and have small somata, and smaller, less complex dendritic arbors. M2 ipRGCs exhibit intermediate characteristics between M4 and M1 cells. Given the key roles of Brn3b in gene regulation associated with ipRGC identity, we next examined whether Brn3b shapes the morphology of ipRGC subtypes. We first compared the morphology of M4 and M2 ipRGCs in Brn3bcKO and control retinas, which both contain the *Opn4^Cre^* allele, by intravitreally injecting a diluted, Cre-dependent AAV2-hSyn-DIO-hM3Gq-mCherry virus. This allowed for sparse mCherry labeling of M4 (identified as mCherry-positive, SMI-32-positive) and M2 (identified as mCherry-positive, SMI-32-negative) ipRGCs (Fig. 3, A and B and fig S7, (*12, 26, 27*)). M1 cells are not labeled at this low titer ((*26*) and see methods). Strikingly, M2 and M4 ipRGCs in Brn3bcKO retinas showed a shift toward morphological features reminiscent of M1 cells, including significant reductions in dendritic arbor diameter, complexity, and total length, as well as soma diameter (Fig. 3, A, D and F). For M4 cells, these changes were most pronounced in the nasal retina, which is the region in control retinas with the largest and most complex M4 cells (fig. S7, (*26*)). Notably, nearly all labeled M2 and M4 cells in Brn3bcKO retinas remain stratified in the ON sublamina (∼10% stratified in the OFF sublamina, not shown), indicating that Brn3b is not required for lamination of M2 and M4 ipRGCs. These data suggest that Brn3b establishes the dendritic and somatic morphology of M2 and M4 ipRGCs that distinguishes them from M1 ipRGCs.

**Fig. 3.**
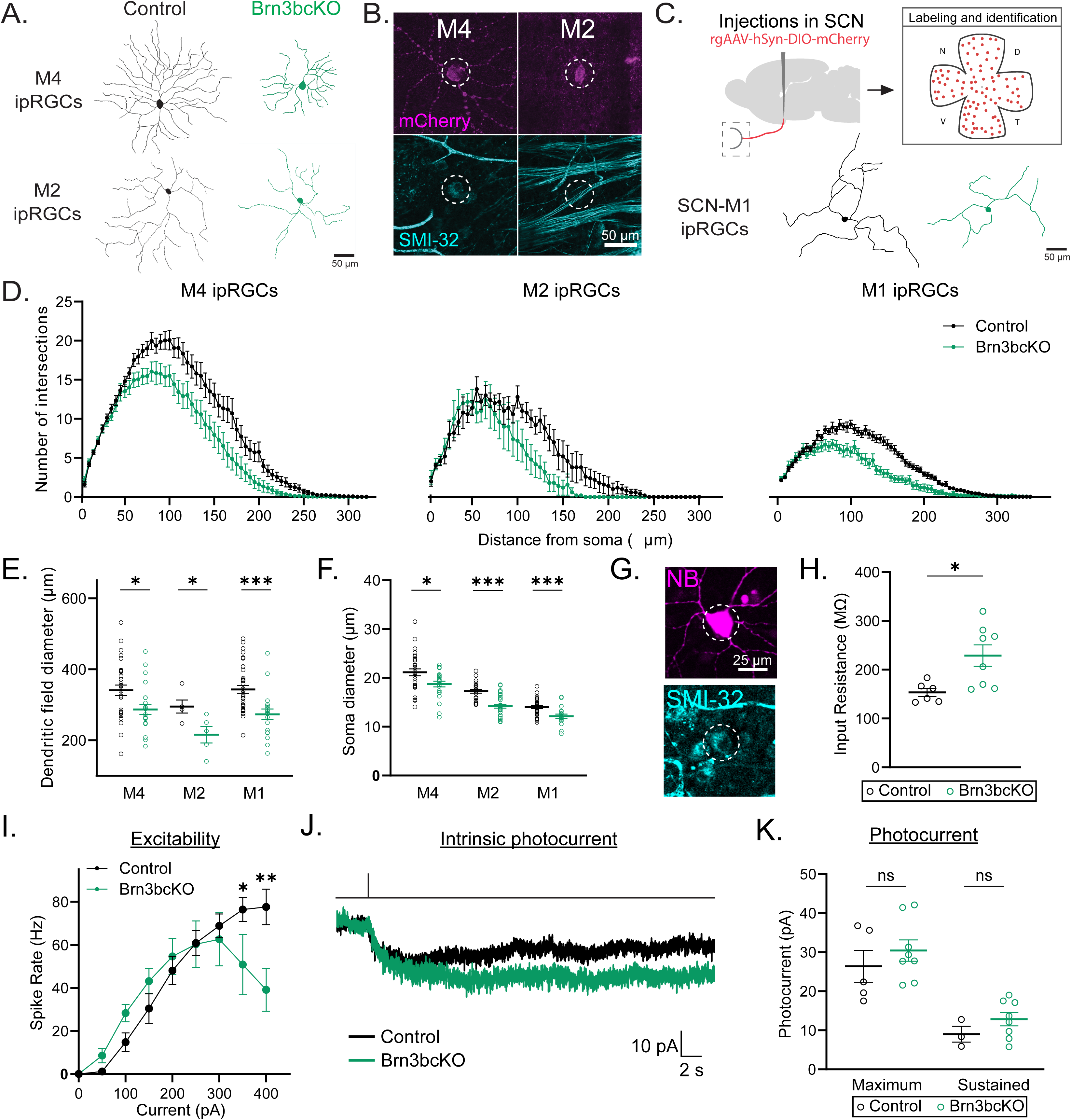
Brn3b tunes morphophysiological properties of ipRGC subtypes. (**A**) Representative dendritic arbor tracing of M4 and M2 ipRGCs in control and Brn3bcKO mice. (**B**) Subtype identified by presence (M4) or absence (M2) of SMI-32 immunolabeling (dashed ellipses), examples are from Brn3bcKO cells. (**C**) (Top) Scheme of retrogradely labeling strategy of M1 ipRGCs. (Bottom) Representative dendritic arbor tracing of M1 ipRGCs in control and Brn3bcKO mice. (**D**) Sholl analysis of M4, M2 and M1 ipRGC subtypes (n = 5-36 cells/group). ipRGCs from Brn3bcKO mice present less complex dendritic arbors compared to control mice. (**E**-**F**) Brn3bcKO showed decreased dendritic field diameter (E) (n = 5-36 cells/group) and soma size (F) (n = 19-36 cells/group) in M4, M2 and M1 ipRGC subtypes than control mice. (**G**) Representative picture of M4 ipRGCs (dashed ellipses) filled with neurobiotin (NB) used for electrophysiological recordings. (**H**) Brn3bcKO M4 ipRGCs showed increased input resistance compared to control M4 ipRGCs (n = 6-8/group). (**I**) M4 ipRGC subtype excitability in control and Brn3bcKO (n =5-7 cells/group) retinas. Brn3bcKO M4 ipRGCs reached peak firing rate by 300pA and then showed marked depolarization block (**J**) Representative traces of photocurrent of M4 ipRGCs from control (black) and Brn3bcKO (*17*) retinas. (**K**) Maximum and sustained phases of photocurrent in M4 ipRGCs (n = 4-6 cells/group). All data are Mean ± SEM, n.s. (not significant) P>0.05, *P<0.05, ***P<0.001, Student’s t- and Mann Whitney U tests and two-way repeated measures ANOVA with Tukey’s multiple comparisons test.

M1 ipRGCs also express Brn3b, albeit at much lower levels than M4 or M2 cells (Fig. 1D)(*16*). Indeed, the suprachiasmatic nucleus-projecting M1 (SCN-M1) ipRGCs reportedly express little to no Brn3b (*16*), suggesting that their morphology may minimally impacted or unaffected in Brn3bcKO animals. To test this, we compared cellular morphology of SCN-M1 ipRGCs in Brn3bcKO versus control retinas. We labeled SCN-M1 ipRGCs using a retrograde, Cre-dependent AAV (rgAAV-hSyn-DIO-hM3Gq-mCherry) injected into the SCN to visualize the dendritic arbor and soma of SCN-M1 cells (Fig. 3C). Contrary to our expectations, we found that SCN-M1 ipRGCs in Brn3bcKO animals have significantly smaller dendritic arbor size, total length, and complexity, as well as significantly smaller soma diameter than control littermates (Fig. 3, C to F). These differences were most pronounced in SCN-M1 ipRGCs of the ventral retina (fig. S8).

These Brn3b-dependent changes in SCN-M1 morphology led us to examine whether SCN-M1 ipRGCs might in fact express Brn3b. We tested this by performing mRNA FISH labeling of Brn3b mRNA in retinas of *Opn4^Cre^* animals where SCN-M1 ipRGCs were retrogradely labeled as described above. Consistent with the observed morphological changes, we detected low levels of Brn3b expression in SCN-M1 ipRGCs (fig. S9). Notably, ventral SCN-M1 cells showed significantly higher Brn3b mRNA labeling than their dorsal counterparts, supporting our observation that ventral SCN-M1 cells of the Brn3bcKO retinas showed more drastic changes in morphology compared to those in the dorsal retina (fig. S9). These findings demonstrate that SCN-M1 ipRGCs express modest amounts of Brn3b, and that even at these lower levels, Brn3b is crucial for regulating both dendritic and somatic morphology. Collectively, our data show that Brn3b expression drives the development of larger somata and larger, more complex dendritic arbors to shape the morphological features that define ipRGC subtype identity.

## Brn3b defines physiological properties of M4 ipRGCs

Intrinsic physiological properties vary across ipRGC subtypes, and shape how features of the visual scene are relayed to downstream areas in the brain. Intriguingly, two key properties, input resistance (M4<M2<M1) and maximum evoked spike frequency (M4>M2>M1) correlate with Brn3b-expression levels across M1, M2, and M4 ipRGCs (fig. S10)(*12, 24*). We therefore tested whether loss of Brn3b would alter these properties in M4 ipRGCs, which show the highest Brn3b expression of all three subtypes. We performed whole cell patch clamp recordings of M4 ipRGCs in Brn3bcKO and control littermates in the presence of synaptic blockers to isolate the intrinsic physiological properties of the cells (Fig. 3G and fig. S11). M4 ipRGCs showed significantly increased input resistance in Brn3bcKO retinas (Fig. 3H). Brn3bcKO M4 cells also reached significantly lower firing rates when injected with positive current compared to those of littermate controls (Fig. 3I and fig. S12). One notable feature of M4 ipRGC firing that further distinguishes it from that of M1 and M2 cells is that even large injections of positive current fail to drive M4 cells into depolarization block (Fig. 3I and fig. S12A)(*12, 24, 27*). However, Brn3bcKO M4 cells did in fact reach depolarization block when injected with positive current, indicating a shift in M4 cell properties toward those of M1 and M2 cells (Fig. 3I and fig. S12B). These differences were still detectable even when normalizing for current density to account for cell size differences, indicating that the spiking properties of M4 cells are affected by loss of Brn3b, and that this phenotype is not secondary to a decrease in cell size (fig. S12). Interestingly, the melanopsin photocurrent of M4 cells was unchanged in amplitude or kinetics, indicating that Brn3b is dispensable for this feature of M4 cells (Fig. 3, J and K; and fig. S13). Overall, our data show that Brn3b regulates both excitability and input resistance of M4 ipRGCs, key features that distinguish this subtype from M1 and M2 cells.

## ipRGC-dependent behaviors are altered in Brn3bcKO animals

The transcriptional and morphophysiological features that define ipRGC subtypes are key to their role in diverse visual behaviors including contrast sensitivity, pupillary light reflex (PLR), and circadian photoentrainment. Given our findings on the critical role of Brn3b in ipRGC subtype transcriptional identity, morphology, and intrinsic physiological properties, we next examined whether ipRGC-dependent behaviors would be altered in Brn3bcKO animals. Contrast sensitivity is crucial for pattern vision and visual perception, and disruption or ablation of non-M1 ipRGCs reduces contrast sensitivity thresholds (*12*). Using the Optomotry system (*28*), we found that Brn3bcKO animals showed significantly decreased contrast sensitivity compared to control littermates, but no change in spatial frequency threshold, indicating that normal contrast sensitivity depends on Brn3b expression in ipRGCs (Fig. 4, A to C and fig. S14).

**Fig. 4.**
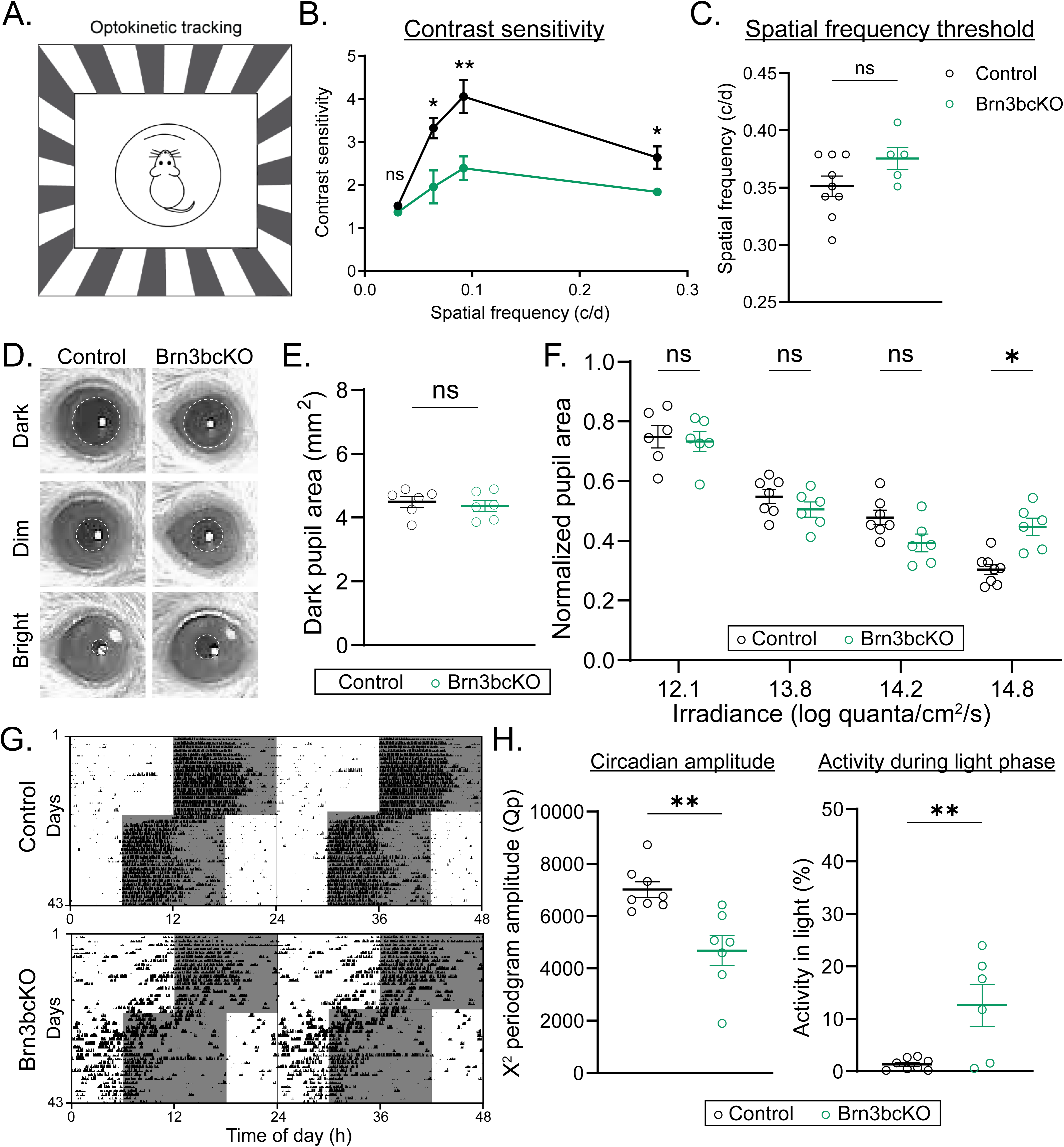
Brn3b is a central regulator of ipRGC-driven behaviors. (**A**) Schematic showing the optokinetic tracking (OKT) behavioral assay. (B) Contrast sensitivity curve at different spatial frequencies. Brn3bcKO mice showed decreased contrast sensitivity compared to control mice (n = 6-11 /group). (C) Spatial frequency threshold measured by OKT at 100% contrast in control and Brn3bcKO (n = 5-9 /group) mice. No differences between Brn3bcKO and control littermates. (D) Representative PLR images from control (left) and Brn3bcKO (right) mice in darkness (top), dim light (middle, 13.8 log quanta cm−2 s−1), and bright light (bottom, 14.8 log quanta cm−2 s−1). Pupils are highlighted with dashed ellipses. (E) Control and Brn3bcKO (n = 6 mice/group) pupil area in the dark. (F) Irradiance-response relationship of PLR in control and Brn3bcKO mice. (G) Representative double-plotted actograms from control (top) and Brn3bcKO (bottom) mice. Mice were initially exposed to a 12:12 LD cycle with 100 lux light during the light phase. Then, the mice were exposed to a 6-hour phase advance. (H) Circadian amplitude measured using the peak amplitude of the χ2 periodogram (left) and percent activity during the light phase (right) in control and Brn3bcKO mice (n = 7-8/group). All data are Mean ± SEM, n.s. (not significant) P>0.05; *P < 0.05; **P < 0.01. Mann-Whitney U test and two-way repeated measures ANOVA with Tukey’s multiple comparisons test.

Pupil constriction is an M1 ipRGC-dependent behavior that regulates the amount of light entering the eye and for improving visual function, allowing animals to see across a broad range of environmental light intensities (*11, 16, 29*). We therefore next tested whether the pupillary light reflex was altered in Brn3bcKO animals by comparing consensual pupil constriction Brn3bcKO and control mice across a range of light intensities at 470 nm (12.1-14.8 log quanta/cm^2^/s) (Fig. 4, D to F). Brn3bcKO animals showed significantly less pupil constriction at the brightest light intensity but no change in kinetics or dark pupil diameter, suggesting that Brn3b-dependent regulation of M1 ipRGC features may be required for normal pupil constriction in bright light (Fig. 4F and fig. S15).

Another behavior regulated by M1 ipRGCs is circadian photoentrainment, the synchronization of an organism’s body clocks to environmental light/dark cycles to properly time activity and physiology rhythms. Misalignment of these rhythms with environmental light/dark cycles has severe consequences for mental and physical health and animal survival (*30, 31*). We assessed whether proper photoentrainment is dependent on Brn3b expression in ipRGCs by tracking voluntary wheel-running activity of littermate Brn3bcKO and control littermates in a 12-hour:12-hour light:dark (LD) cycle at 100 lux. All mice photoentrained with a similar period length and total daily activity (fig. S16). However, we observed significantly lower circadian amplitudes in Brn3bcKO mice, indicating that loss of Brn3b in ipRGCs dampens photoentrainment (Fig. 4, G and H; fig. S16 and fig.17). Additionally, Brn3bcKO mice showed significantly higher activity during the light phase of the LD cycle and more variable, phase-advanced, activity onset, suggesting impaired light signaling to the SCN (Fig. 4H and fig. S16). These data indicate that Brn3b expression in SCN-M1 ipRGCs is required for their proper function in circadian photoentrainment. Together, our findings unveil the critical roles of Brn3b in the genetic and morphophysiological tuning of ipRGC subtypes that is critical in shaping their proper function across multiple visual behaviors.

## Discussion

A unique blend of molecular and cellular features enables each ipRGC subtype to precisely regulate downstream brain circuits and behaviors that operate over a range of timescales and environmental contexts. The transcriptional programs underpinning this neuronal diversity were previously unknown. Here, we identify graded expression of the transcription factor Brn3b across ipRGC subtypes as a key mechanism in shaping their genetic and morphophysiological features. Higher levels of Brn3b expression in M4 ipRGCs differentiate these cells from M1 and M2 ipRGCs, and knockout of Brn3b shifts the properties of M4 cells towards those of M1 and M2 cells. The cellular changes in ipRGCs following Brn3b knockout result in deficits in several visual behaviors, including in contrast sensitivity, a behavior linked to M4 ipRGCs (*16*). Surprisingly, we find that even SCN-M1 ipRGCs show low levels of Brn3b expression that are critical for shaping cellular properties and their role in circadian photoentrainment. SCN-M1 cells are spared in mouse lines where diphtheria toxin expression from the Brn3b promoter is driven in ipRGCs, which had resulted in the conclusion that these cells did not express Brn3b. However, our data suggest that SCN-M1 survival in this line is simply due to low expression of Brn3b and therefore diphtheria toxin and that even this low Brn3b expression shapes the features and resulting behavioral functions of SCN-M1 cells.

Our findings also suggest that ipRGC subtype identity is shaped by multiple transcriptional programs, many of which are under the control of Brn3b. It will be interesting to determine how ipRGC features that develop independently of Brn3b activity, including the laminar positioning of ipRGCs, are genetically regulated. Intriguingly, Brn3b continues to modify ipRGC identity from early development through adulthood, partly by fine-tuning melanopsin expression. Whether extrinsic cellular signals and experiences in animals may control Brn3b activity to regulate the cellular functions of ipRGCs even into adulthood remains to be elucidated. Together, our study opens new windows into the genetic tuning of retinal cell types and paves the way for future studies on how molecular programs functionally shape cellular identity in the brain.

## Materials and Methods

### Animals

All procedures were approved by the Animal Care and Use Committee at Northwestern University. Male and female mice were used for all experiments at embryonic (E13.5-E15) and adult (P40-P120) stages. For RNAscope Fluorescence *in situ* hybridization and immunohistochemical experiments we utilized *WT*, *Opn4^Cre/+^* (*13*), RRID:IMSR_JAX:035925), *Opn4^Cre/+^; Brn3b^cKOAP^* (*15*), RRID: IMSR_JAX:010559) and *Opn4^CreERT2/+^; Brn3b^cKOAP^* mice (*16*), RRID:IMSR_JAX:035926). For morphological and physiological experiments, we used *Opn4^Cre/+^* and *Opn4^Cre/+^; Brn3b^cKOAP^* mice. TRAPseq experiments were conducted using *Opn4^Cre/+^ Rpl22^HA^* (*18*), RRID: IMSR_JAX:029977) and *Opn4^Cre/+^; Brn3b^cKOAP^ Rpl22^HA^* mice. For behavioral experiments we utilized *Opn4^Cre/+^* and *Opn4^Cre/+^; Brn3b^cKOAP^*mice.

### TRAPseq

Eyes from postnatal day 60 (P60) mice expressing HA-Rpl22 in ipRGCs or control mice were enucleated, and eyecups were dissected in cold nuclease-free 1X PBS with 100 μg/ml cycloheximide. Retinas from the same animal were combined and homogenized in lysis buffer (50 mM Tris-HCl pH 7.4, 100 mM KCl, 12 mM MgCl_2_, 1% NP-40, 0.4 unit/ml RNase inhibitor (Promega, N2511), 1 mM DTT, 100 μg/ml cycloheximide) with a dounce homogenizer. Lysates were incubated with 1 µg/µl heparin (Sigma, H3393) for 2 minutes on ice prior to centrifugation. The lysates were spun down at 10,000*g* for 10 minutes at 4C and the supernatant was used for immunoprecipitation using an antibody against HA-tag (Sigma, H3663) with Dynabeads protein G (Invitrogen) for overnight at 4C. 1.5% of lysate was saved as an input. After extensive washes with cold washing buffer (50 mM Tris-HCl pH 7.4, 300 mM KCl, 12 mM MgCl_2_, 1% NP-40, 0.5 mM DTT, 100 μg/ml cycloheximide), immunoprecipitated RNA and input RNA were purified with a RNeasy Micro Kit (Qiagen) following the manufacturer’s instructions and eluted in 10 µl of water. 2 µl of purified RNA were used for sequencing library preparation using the Smart-seq3 protocol (*32*) with modifications. After reverse transcription followed by PCR amplification, 25 ng of cDNA was used for the Tn5 tagmentation reaction. Tagmentated cDNA was amplified with 8 cycles of PCR using primers containing indexes and Illumina adaptors. The RNA libraries were quantified by Qubit (Thermo Fischer Scientific) and bioanalyzer (Agilent) and sequenced on the Illumina NextSeq 550 platform to obtain 37 bp paired-end reads. Two to four biological replicates were performed for all conditions.

### RNA-seq analyses

For TRAP-seq analyses, Smart-seq3 prepared libraries were aligned to the mm10 reference genome using the Smart-seq3 github pipeline (https://github.com/sandberg-lab/Smart-seq3) (*32*). Transcripts considered for analyses were selected based on higher normalized counts following HA immunoprecipitation in HA-Rpl22 expressing retina compared to HA immunoprecipitation in control retina. Differential gene expression analyses were performed using pair-wise negative binomial tests with edgeR (*33*) and the false discovery rate (FDR) was calculated for all genes.

The top 100 genes enriched or de-enriched in ipRGCs following HA-Rpl22 immunoprecipitation compared to total retinal RNA were used to generate modules for visualization on the scRNA-seq UMAP plot using Seurat 4(*34*). Genes upregulated or downregulated in Brn3b cKO ipRGCs compared to control ipRGCs (log_2_ fold change >1 or <-1) or not changing (absolute log_2_ fold change < 0.05) following HA-Rpl22 immunoprecipitation were intersected with the top 2000 genes with the highest standardized variance in ipRGCs calculated using Seurat 4(*34*). The fraction overlap was derived using the number of overlapping genes divided by the number of Brn3bcKO-upregulated, downregulated, or not changing genes. For comparisons of selected transcripts using HA-Rpl22 immunoprecipitation of Brn3bcKO and control retinas, we considered the dominant HA-labeled cell types including ipRGCs and a small group of cone cells. To adjust for changes in the ratio of these cells upon knockout of Brn3b, the transcript FC of ipRGC-enriched genes was normalized by the FC in number of HA-labeled ipRGCs relative to the number of HA-labeled cones.

scRNA-seq datasets from RGCs in adult mice with cell type and subtype annotations were obtained (*3*) and analyzed using Seurat 4(*34*). Briefly, all RGCs or ipRGCs were extracted for downstream analyses. Read counts were normalized and integrated across three biological replicates using fastMNN followed by UMAP embedding (*35*). Module scores were calculated from the average expression of genes within the module, subtracted by the levels of similarly expressed control genes (*34, 36*). To identify genes expressed in a graded pattern across M1 to M6 ipRGCs, differentially expressed genes in clusters containing M1 and M2/3 ipRGCs were compared with those in clusters containing M4, M5, and M6 ipRGCs and an unidentified group (cluster 22) using the Wilcoxon rank sum test. Genes enriched in M4-M6 ipRGCs and cluster 22, including Brn3b, were denoted as Brn3b^High^. Genes enriched in M1 and M2/M3 ipRGCs were denoted as Brn3b^Low^.

### Viral infection

For morphological studies we used two strategies to label different ipRGC subtypes. To sparsely label M4 and M2 ipRGCs (*26*), *Opn4^Cre/+^* and *Opn4^Cre/+^; Brn3b^cKOAP^* mice between P40-60 were anesthetized by intraperitoneal (IP) injection of Avertin and a 30-gauge needle was used to open a hole in the *ora serrata*. Each eye was intravitreally injected with 1 μL of AAV2-hSyn-DIO-hM3Gq-mCherry (∼8 x 10^11^ GC/mL, Addgene) using a custom Hamilton syringe (Borghuis Instruments) with a 33-gauge needle (Hamilton). To label M1 ipRGCs, mice were anesthetized with isoflurane (4% for induction in chamber and 2% with mask, Kent Scientific VetFlo system) and then bilaterally injected with 150 nl of rgAAV-hSyn-DIO-hM3Gq-mCherry (∼8 x 10^12^ GC/mL, cat. #: 44361-AAVrg, Addgene) in the suprachiasmatic nucleus (SCN) (AP: -0.2, ML: ±0.15, DV: 5.85) using a stereotaxic injector (Neurostar) controlled by the software Stereodrive at an injection rate of 30 nl/min.

To label ipRGCs for electrophysiological studies *Opn4^Cre/+^*and *Opn4^Cre/+^; Brn3b^cKOAP^* mice between P40-60 were anesthetized by IP injection of Avertin and a 30-gauge needle was used to open a hole in the *ora serrata* as described above. Each eye was intravitreally injected with 1 μL of AAV2-hSyn-DIO-mCherry (∼8 x 10^12^ GC/mL, cat. #: 50459-AAV2, Addgene) using a custom Hamilton syringe (Borghuis Instruments) with a 33-gauge needle (Hamilton). After all surgery procedures, mice were subcutaneously injected with 2 mg/kg of meloxicam (Covetrus).

### Tamoxifen injection

Tamoxifen (TMX, Sigma-Aldrich) was dissolved in corn oil at a concentration of 20 mg/ml by shaking overnight at 37°C and then the solution was stored at 4°C for the duration of injections. *Opn4^CreERT2/+^; Brn3b^cKOAP^* mice at P60 were IP injected once every 24 hours for a total of 5 consecutive days with 100µl of tamoxifen or Vehicle (corn oil) (*16*). Mice were closely monitored and no adverse reactions to the treatment were noticed. Mice were euthanized 60 days after the first injection for RNA fluorescence *in situ* hybridization and immunohistochemical studies.

### RNA fluorescence *in situ* hybridization (FISH)

To obtain tissue sections, mice were anesthetized by IP injection of Avertin and sacrificed by cervical dislocation. Eyes were then enucleated, and eyecups were dissected in nuclease-free PBS and fixed eyecups in 4% paraformaldehyde (PFA) solution in PBS for 24 hours at 4°C (*37*). A large relieving cut was made in the nasal margin of the eyecup. For experiments at embryonic stages, WT pregnant females were anesthetized by IP injection of Avertin and sacrificed by cervical dislocation. Embryos were dissected, fixed in PFA 4% overnight. Eyecups and embryos were then washed in PBS and cryoprotected in a 30% sucrose solution in PBS overnight at 4°C. Eyecups and embryos were then embedded and frozen in OCT using dry ice. 20µm sections were obtained on a Leica CM1950 cryostat and mounted directly onto SuperFrost Plus slides (Fisher). The tissue was processed according to the RNAscope Multiplex Fluorescent v2 assay (Advanced Cell Diagnostics) instructions provided by the manufacturer. Probes (Advanced Cell Diagnostics) and fluorophores (TSA Plus Fluorescence kits, Akoya Biosciences) are listed in Table S1. The tissue was then incubated in DAPI solution (Sigma) prepared in nuclease-free PBS for 10 minutes and mounted using ProLong Glass Antifade Mountant (Thermo).

Sections were imaged on a Leica SP5 confocal microscope. For quantification, high magnification images (367.38 µm x 367.38 µm with a pixel size of 0.71 µm) with a z-stack size of 1 µm were taken. Because ipRGCs are a sparse population of RGC, regions of interest (ROIs) around Opn4 mRNA or mCherry mRNA (Fig. 2B, fig. S3, fig. S4, fig. S6 and fig. S9) were manually drawn. The mean pixel intensity, per ROI, per stack, per probe was established using ImageJ. Analyses were performed on 336 cells of 6 retinas from *Opn4^Cre/+^* mice and 213 cells of 6 retinas from *Opn4^Cre/+^; Brn3b^cKOAP^* mice (Fig. 2B), 371 cells of 2 retinas from Vehicle-injected *Opn4^CerERT2/+^ Brn3b^cKOAP^* mice and 288 cells of 2 retinas from TMX-injected *Opn4^CerERT2/+^ Brn3b^cKOAP^* mice (Fig. 2C and fig. S6), 24 cells of 2 retinas from *Opn4^Cre/+^* mice (fig. S9). Qualitative analysis was done in 18 retinas from 9 WT embryos (fig. S2).

To perform RNA FISH in whole mount retinas, mice were anesthetized by IP injection of Avertin and sacrificed by cervical dislocation. Eyes were then enucleated, and retinas were dissected in nuclease-free PBS and eyecups were fixed in 4% paraformaldehyde solution in PBS overnight at 4°C. The nasal margin of the retina was labeled with a cut for orientation. Retinas were then washed in PBS, followed by dehydration in a graded methanol (MeOH)/PBS series (50%, 75% MeOH-PBS, 100% MeOH) with agitation at room temperature for 5 min each. Retinas were stored at 100% MeOH at -20 C overnight. The tissue was rehydrated in a reverse MeOH/PBS series and washed three times for 5 min each in PBS. Retinas were incubated with RNAscope Protease Plus Reagent (Advanced Cell Diagnostics) for 30 min at 40 C. Retinas were washed and incubated with probes (Table S1) overnight. Retinas were then washed and post-fixated for 10 min at room temperature in PFA 4%, washed and processed according to the RNAscope Multiplex Fluorescent v2 assay (Advanced Cell Diagnostics) instructions provided by the manufacturer. The tissue was mounted and sealed using ProLong Glass Antifade Mountant (Thermo Fisher Scientific).

Whole mount retinas were imaged on a Leica SP5 confocal microscope. For quantification, high magnification images (183.69 µm x 183.69 µm with a pixel size of 0.36 µm) with a z-stack size of 1 µm were taken. ROIs around Opn4 mRNA and control background were manually drawn and the mean of pixel intensity, per ROI, per stack, per probe was established using ImageJ. Images were processed, the mean background was subtracted and label intensity data for each ROI on every whole mount retina was normalized against the maximum intensity value observed within that retina. Analyses were performed 182 cells of 2 retinas from *Opn4^Cre/+^* mice and 241 cells from 2 retinas from *Opn4^Cre/+^; Brn3b^cKOAP^* mice (Fig. 1, I to K and fig. S4). Zcchc12-positive (Zcchc12^+^) cells were considered when mean intensity of ROIs were positive, while Zcchc12-negative (Zcchc12-) cells were computed when ROIs presented mean intensity were 0 or negative.

### Immunohistochemical procedures

Mice were anesthetized by IP injection of Avertin and sacrificed by cervical dislocation. Eyes were then enucleated, and retinas were dissected in PBS. A large relieving cut was made in the nasal margin of the eyecup prior to removing the retina for experiments in which retinal orientation was tracked. Retinas were fixed in PFA 4% in PBS for 30-60 minutes at RT and washed in PBS (3 x 10 minutes). Retinas were then blocked at 4 °C overnight in 6% normal goat serum in 0.3% Triton PBS prior to incubating in primary antibody solution for 2-3 nights at 4 °C (Table S2). Then, retinas were washed in PBS (3 x 10 minutes) at RT and incubated in secondary antibody solution for 2 hours at RT (Table S2). Retinas were washed and mounted using Fluoromount (Sigma). Primary and secondary antibody solutions were made in 3% normal goat serum in 0.3% Triton PBS.

Whole mount retinas were imaged on a Leica SP5 confocal microscope. For quantification, high magnification images (183.69 µm x 183.69 µm with a pixel size of 0.36 µm) with a z-stack size of 1 µm were taken and processed using ImageJ. Total number of cells in the retinal ganglion cell layer (GCL) and the density of cells of the outer nuclear layer (ONL) were quantified in 6 retinas from *Opn4^Cre/+^ Rpl22^HA^* mice and 5 retinas from *Opn4^Cre/+^; Brn3b^cKOAP^ Rpl22^HA^* mice (fig. S18). ROIs around melanopsin-immunolabel and control background were manually drawn and the mean pixel intensity, per ROI, per stack, per probe was established using ImageJ. Images were processed, the mean background was subtracted and label intensity data for each ROI on every whole mount retina was normalized against the maximum intensity value observed within that retina. Analyses of melanopsin levels were performed in 600 cells of 5 retinas from *Opn4^Cre/+^* mice (Fig. 2, D to F and fig. S5), 600 cells from 4 retinas from *Opn4^Cre/+^; Brn3b^cKOAP^* mice (Fig. 2, D to F and fig. S5), 800 cells of 4 retinas from Vehicle-injected *Opn4^CreERT2/+^; Brn3b^cKOAP^* mice (Fig. 2, G and H), 761 cells from 4 retinas from TMX-injected *Opn4^Cre/+^; Brn3b^cKOAP^*mice (Fig. 2, G and H). For morphological studies in sparse and retrogradely labeled ipRGCs, dendritic arbor tracings were performed using the Simple Neurite Tracer toolbox (https://github.com/morphonets/SNT) for ImageJ in the mCherry channel (Fig. 3, B and C; fig. S7 and fig. S8). ChAT immunolabeling allowed determination of the dendritic stratification level in ON or OFF sublayers of the inner plexiform layer (IPL). Using the intravitreal sparse labeling approach, M4 ipRGCs were identified as mCherry-positive, ON-stratifying, SMI-32-positive (Fig 3B); while M2 ipRGCs were identified as mCherry-positive, ON-stratifying, SMI-32-negative positive (Fig 3B). All retrogradely labeled cells from SCN were identified as M1 ipRGCs (mCherry-positive, OFF-stratifying, SMI-32-negative). All morphological parameters were calculated using the Simple Neurite Tracer toolbox.

### *Ex vivo* retina preparation for electrophysiology

Mice were eye injected with AAV2-hSyn-DIO-mCherry 1-2 weeks prior to electrophysiology recordings to permit targeted recordings of ipRGCs (fig. S11). 60-90 day old mice were dark-adapted for at least one hour and sacrificed by CO_2_ asphyxiation. Eyes were enucleated and the retinas were dissected under dim red light in carbonated (95% O_2_-5% CO_2_) Ames’ medium (Sigma-Aldrich). Retinas were sliced in half along the nasal-temporal axis and incubated in carbonated Ames’ medium in the dark at 26 °C for at least 1 hour prior to use. Retinas were mounted on a glass-bottom recording chamber and anchored using a platinum ring with nylon mesh (Warner Instruments). The chamber was placed on an electrophysiology rig and the tissue was perfused with Ames’ medium with synaptic blockers. All recordings were made at 25-26 °C.

### Solutions for electrophysiology

All recordings were made in Ames’ medium with 23 mM sodium bicarbonate. Synaptic transmission was blocked with 100 μm DNQX (Tocris), 20 μm L-AP4 (Tocris), 100 μm picrotoxin (Sigma-Aldrich), and 20 μm strychnine (Sigma-Aldrich) in Ames’ medium. All recordings were made with internal solution containing 125 mM K-gluconate, 2 mM CaCl_2_, 2 mM MgCl_2_, 10 mM EGTA, 10 mM HEPES, 10 mM Na_2_-ATP, 0.5 mM Na-GTP, and 0.3% Neurobiotin (Vector Laboratories).

### ipRGC electrophysiological recordings and data analysis

Recordings were made using borosilicate pipettes (Sutter Instruments) with resistances between 4-6 MΩ. Data was collected using a Multiclamp 700B amplifier (Molecular Devices) with pClamp 10 acquisition software. Reported voltages are corrected for 14 mV liquid junction potential between the Ames’ medium and K-gluconate internal solution. To identify M4 ipRGCs, mCherry-expressing cells were visualized with epifluorescence and M4 ipRGCs were targeted based on soma size. A blue LED light (∼480 nm) was used to deliver light stimuli to the retina through a ×60 water-immersion objective. The photon flux was attenuated using neutral density filters (Thor Labs).

To measure input resistance, cells were voltage clamped at –70 mV and hyperpolarized with a – 10 mV step. Input resistance was calculated with Ohm’s law using the steady state current during the voltage step. To measure the excitability, cells were held at ∼-70 mV in current clamp and then 0.5 second current injections ramping from 50 pA to 400 pA were applied.

To measure the intrinsic light response, cells were voltage clamped at –70 mV and the photocurrent was recorded following a 50 ms full-field flash of bright light with an intensity of 6.08 × 10^15^ photons · cm^−2^ · s^−1^. Only one light response was recorded per retina piece to ensure that light adaptation was consistent across recordings. The Maximum photocurrent represents the greatest change from baseline after light onset and the Sustained photocurrent was calculated as the average change from baseline during the 40-50 seconds after light onset. The Early, Intermediate, and Late timepoints were calculated as defined in (*38*) with the following timeframes after light onset: Early (141.7 - 440.4 ms), Intermediate (2857.7 - 6598.2 ms), and Late (9062.3 -14062.3 ms). Photocurrent values were reported as the absolute value of the current for each component. Electrophysiological data was analyzed in Python using the PyABF package (https://pypi.org/project/pyabf/) and custom scripts. The code is available on GitHub.

To ensure recorded neurons were M4 ipRGCs, tissue was fixed overnight in 4% paraformaldehyde solution in PBS at 4°C. Retinas were then washed in PBS for 45 minutes and blocked overnight in 6% normal donkey serum in 0.3% Triton-X. Retinas were then incubated in primary antibody solution for 3-5 nights at 4 °C (Table S2), washed for 45 minutes (3 x 15 minutes) at room temperature, and incubated in secondary antibody solution overnight at 4 °C (Table S2). Sections were then washed in PBS and mounted on glass slides. Cells were defined as M4 ipRGCs based on the expression of SMI32 and the presence of an intrinsic light response.

### Optokinetic tracking response

Optokinetic tracking response experiments were performed between Zeitgeber times (ZT) 6 and 8. Spatial frequency thresholds were assessed using the virtual OptoMotry system (Cerebral Mechanics, Lethbridge, Alberta) as described previously (*12, 28, 37*). A vertical sine wave grating was projected as a virtual cylinder in 3D coordinate space on computer monitors arranged in a quadrangle around a testing arena. Unrestrained animals were placed individually on an elevated platform at the epicenter of the arena. Experimenters were blind to the genotype of all animals. The experimenter used a video image of the arena from above to view the animal and followed the position of its head in real time to determine when the mouse “tracked” which was defined as when the head and neck movements of the animals moved in the same direction as the drifting grating. Animals were presented with stimuli of increasing spatial frequencies at 100% contrast to assess the spatial frequency threshold of the animal in both clockwise and counterclockwise directions. Because the temporal-to-nasal stimulus movement in the visual field evokes tracking, clockwise and counterclockwise stimulation can be used to assess the spatial frequency threshold of each eye individually (*39*). All measurements were conducted using a cylinder rotation of 12°/sec, and these thresholds were measured for each eye and averaged for each mouse.

Contrast thresholds were identified using similar procedures, except that testing began with a grating of maximum (∼100%) contrast, which was then systematically changed until the minimum contrast to elicit tracking was identified. Thresholds were measured at 4 spatial frequencies (0.031, 0.064, 0.092, 0.272 c/d), and a contrast sensitivity function was generated by calculating a Michelson contrast using the screen luminance (max-min)/(max+min). A complete set of spatial frequency and contrast thresholds for an animal through each eye were generated in 10-15 minutes. Experimenters were blind to an animal’s experimental condition and previously recorded thresholds, and adjunct observers were used to validate threshold assessments.

### Voluntary wheel running behavior

Wheel running activity was recorded by individually housing P90 male mice in cages with a running wheel. Activity was recorded using ClockLab Data Collection software (Actimetrics). Mice were first exposed to a 12:12 light dark (LD) cycle with 100 lux light during the light phase for 3 weeks. Then, animals were exposed to a 6-hour phase advance. Mice that stopped running during the experiment were excluded. All data analysis was performed using ClockLab 6 analysis software (Actimetrics). Circadian amplitude, total activity, percent activity in light and onset error were measured in the 8 days preceding phase advance.

### Pupillometry

For pupillary light reflex (PLR), experiments were performed between ZT 6-8. Experimenters were blind to the genotype of all animals. Consensual PLR was measured by delivering a 470nm light stimulus to one eye while simultaneously recording the other eye using a Sony Handycam camcorder. An LED light source was band passed using a 470nm bandpass filter (Thorlabs) and light intensity was attenuated using neutral density filters (Thorlabs). Mice were dark-adapted for at least 30 minutes prior to any light exposure and were then manually restrained by hand. A 2-5 second baseline recording was taken prior to delivering the 5 seconds light stimulus. The order in which animals were tested after dark adaptation was randomized. Maximum pupil area was quantified *post hoc* using the oval tool in ImageJ. Steady-state pupil area was then calculated by averaging the pupil area during the last 3 seconds of the stimulus. In analyses looking at the time course of PLR, videos were analyzed using DeepLabCut and data were fitted using a single exponential decay function to measure the time constant tau.

### Statistical comparisons

All graphs and statistical analyses were performed using Graph Pad Prism 10.1.2 (RRID: SCR_002798). Normal distribution of data was tested using Shapiro-Wilk test. When comparing two groups, Student’s t- or Mann Whitney U tests were used. For multiple statistical comparisons, we performed two-way ANOVA with repeated measures followed by Tukey’s *post hoc* test. Significance was concluded when P < 0.05.

**Table S1.** List of mRNA probes and fluorophores used in this study. All fluorophores were used 1:1000.

| **Probe** | **Catalog number** | **Tissue** | **Fluorophore** |
| --- | --- | --- | --- |
| Mm-RBPMS | 527231 | Retinal sections | Fluorescein (NEL741001KT) |
| Mm-Opn4-C2 | 438061-C2 | Retinal sections | Cyanine 3 (NEL744001KT) |
| Mm-Pou4f2-C3 | 558481-C3 | Retinal sections | Cyanine 5 (NEL745001KT) |
| Mm-Opn4 | 438061 | Retinal sections | Cyanine 3 (NEL744001KT) |
| Mm-Eomes-C2 | 429641-C2 | Retinal sections | Fluorescein (NEL741001KT) |
| Mm-mCherry-C2 | 431201-C2 | Retinal sections | Cyanine 3 (NEL744001KT) |
| Mm-Opn4 | 438061 | Whole mount retinas | Fluorescein (NEL741001KT) |
| Mm-Chrna6-C2 | 467711-C2 | Whole mount retinas | Cyanine 3 (NEL744001KT) |
| Mm-Zcchc12-C3 | 524441-C3 | Whole mount retinas | Cyanine 5 (NEL745001KT) |

**Table S2.** List of antibodies and tracers used in this study. The dilution of all secondary antibodies and streptavidin was 1:500.

| Purpose | Primary Ab / Tracer | Dilution | Secondary Ab |
| --- | --- | --- | --- |
| To characterize *Opn4<sup>Cre/+</sup>* *Rpl22<sup>HA</sup>* and *Opn4<sup>Cre/+</sup>*; *Brn3b<sup>cKOAP</sup>* *Rpl22<sup>HA</sup>* lines (fig. S18) | Rabbit anti-HA (Abcam, ab9110) | 1:500 | Donkey anti-rabbit Alexa 488 (Thermofisher, A-21206) |
| To test the role of Brn3b on melanopsin expression (Fig. 2) | Rabbit anti-melanopsin (ATS, N38) | 1:1000 | Donkey anti-rabbit Alexa 488 (Thermofisher, A-21206) |
| Morphological studies (sparse labeling) (Fig. 3) | Rabbit dsRed (Takara, 632496) | 1:500 | Donkey anti-rabbit Alexa 594 (ThermoFisher, A-21207) |
|  | Goat anti-ChAT (Sigma, AB144P) | 1:250 | Donkey anti-goat Alexa 488 (Invitrogen, A11055) |
|  | Mouse anti-SMI32 (Biolegend, 801701) | 1:500 | Donkey anti-mouse Alexa 647 (Invitrogen, A31571) |
| Morphological studies (filled cells) (Fig. 3) | Neurobiotin (VectorLabs, SP-1120-50) | - | Streptavidin 546 (Invitrogen, S11225) |
|  | Chicken anti-GFP (Abcam, ab13970) | 1:500 | Donkey anti-chicken Alexa 488 (Jackson IRL, 703-545-155) |
|  | Goat anti-Brn3b (Abcam, ab56026) | 1:250 | Donkey anti-goat Alexa 647 (Invitrogen, A21447) |
| Electrophysiological studies (filled cells) (Fig. 3; fig. S11 and fig. 13) | Neurobiotin (VectorLabs, SP-1120-50) | - | Streptavidin 546 (Invitrogen, S11225) |
|  | Mouse anti-SMI32 (Biolegend, 801701) | 1:500 | Donkey anti-mouse Alexa 488 (Abcam, ab175472) |

## Acknowledgments

We thank Dr. Samer Hattar for the gift of *Opn4^Cre/+^* and *Opn4^CreERT2/+^*mice.

## Author contributions

Conceptualization: MLA, YY and TMS.

Methodology: MLA, JDB, OAPP, SKL and TY.

Investigation: MLA, JDB, OAPP, TY, YY and TMS.

Visualization: MLA, JDB, TY, YY and TMS.

Funding acquisition: YY and TMS.

Project administration: TMS.

Supervision: YY and TMS.

Writing – original draft: MLA and TMS.

Writing – review & editing: MLA, JDB, OAPP, TY, YY and TMS.

## Supplemantary

**Figure S1. Profiling the ipRGC transcriptome using TRAP.** (**A**) ipRGCs identified from publicly available scRNA-Seq profiles of RGCs (*3*), re-analyzed using dimensionality reduction, and visualized with UMAP. Established genetic markers are used to annotate ipRGC subtypes. (**B**) Immunoprecipitation of HA-Rpl22 expressed selectively in *Opn4^Cre/+^* retinal cells followed by sequencing of ribosome-bound mRNA and total retinal mRNA reveals enrichment and de-enrichment of thousands of transcripts as indicated in red or blue, respectively (n = 4 biological replicates, log2FC>1, FDR<0.05). (**C**) Top 100 genes isolated from *Opn4^Cre/+^ ; Rpl22^HA^* retinal cells that are enriched in ipRGCs, among all RGC cell types. (**D**) Brn3b, Opn4, Chrna6 and Zcchc12 expression in ipRGC clusters.

**Figure S2. Opn4 is expressed after RGC specification.** RNAscope fluorescent in situ hybridization (FISH) labeling in developing eyecups for Brn3b (cyan), RBPMS (green), and Opn4 (Magenta) mRNA at various embryonic stages. Opn4 expression is undetectable until E15.5, after RGC specification.

**Figure S3. Brn3bcKO mice present decreased Brn3b mRNA expression in ipRGCs.** (**A**) Schematic representation RNAscope in cross-sectioned retinas. (**B)** Brn3b mRNA levels are significantly decreased in ipRGCs of Brn3bcKO mice compared to control (n = 216-336 cells/group) littermates. Lines are median values, ****P<0.001, Mann Whitney U test.

**Figure S4. Brn3bcKO mice present increased Opn4 mRNA expression in ipRGCs**. (**A**) Schematic representation RNAscope in flat-mount retinas. (**B)** Representative images of Opn4 mRNA labeling in flat mount retinas. (**C**) Opn4 mRNA levels are significantly increased in ipRGCs of Brn3bcKO mice compared to control littermates (n = 182-241 cells/group). Lines are median values, ***P<0.001, Mann Whitney U test.

**Figure S5. Opn4 protein levels in flat mount retinas.** (**A**) Schematic representation Opn4 immunolabeling in flat-mount retinas. (**B**) Representative quantification of total number of melanopsin-positive cells (each dot represents a cell) in control and Brn3bcKO retinas. (**C**) No differences in the total number of melanopsin-positive cells were observed between control and Brn3bcKO (n = 4-5 retinas/group) mice. Lines are median (B-C), n.s. (not significant) P>0.05, Mann Whitney U test.

**Figure S6. Characterization of inducible-Brn3bcKO (iBrn3bcKO) mice.** (**A**) Schematic representation of RBPMS, Brn3b and Opn4 expression during development. (**B**) Schematic protocol to induce Cre-recombinase expression in iBrn3bcKO mice. (**C**) Representative images of RNAscope labeling using Brn3b and Opn4 mRNA probes in retinal sections from Vehicle- or Tamoxifen (TMX)-injected iBrn3bcKO mice. Dashed ellipses show regions of interest of ipRGC. (**D**) Brn3b mRNA levels are significantly decreased in ipRGCs from TMX-injected iBrn3bcKO compared to control mice (n = 288-371 cells/group).

**Figure S7. Nasal M4 ipRGCs show the most pronounced changes in Brn3bcKO retinas.** (**A**) Schematic of the ipRGC sparse labeling protocol. (**B**) Representative traces of nasal and temporal M4 ipRGCs in control and Brn3bcKO retinas. (**C-D**) Sholl analysis (top), total dendritic length and dendritic field diameter (bottom) from nasal (C) and temporal (D) Brn3bcKO and control retinas (n = 9 – 18 cells/group). All data are Mean ± SEM, n.s. (not significant) P>0.05, *P<0.05, **P<0.01, Student’s t-test.

**Figure S8. Ventral M1 ipRGCs show the most pronounced changes in Brn3bcKO retinas.** (**A**) Schematic of retrograde labeling of SCN-projecting ipRGCs and morphological study procedure. (**B**) Representative traces of dorsal and ventral M1 ipRGCs in control and Brn3bcKO retinas. (**C-D**) Sholl analysis (top), total dendritic length and dendritic field diameter (bottom) from nasal (C) and temporal (D) Brn3bcKO and control retinas (n = 8 – 21 cells/group). All data are Mean ± SEM, n.s. (not significant) P>0.05, *P<0.05, **P<0.01, Student’s t-test.

**Figure S9. Ventral SCN-M1 ipRGCs show higher levels of Brn3b mRNA than ventral SCN-M1 ipRGCs.** (**A**) Schematic of retrograde labeling of SCN-projecting ipRGCs and following RNAscope experiments. (**B**) Representative images of RNAscope labeling using mCherry and Brn3b probes in retinal sections from retrogradely labeled retinas from the SCN. Dashed ellipses show regions of interest of mCherry-positive cells (i.e. SCN-ipRGCs). (**C**) Brn3b mRNA intensity quantification in dorsal and ventral SCN-ipRGCs (n = 9-15 cells/group). Data are Mean ± SEM, *P<0.05, Mann Whitney U test.

**Figure S10. Brn3b expression correlates with multiple features of ipRGC subtypes.** Schematic representation of correlation between Brn3b expression levels (blue) with melanopsin expression levels (red); morphological (orange) and physiological properties (green).

**Figure S11. Schematic of electrophysiological recordings of M4 ipRGCs from control and Brn3bcKO mice.** Timeline of intravitreal injections to label ipRGCs, electrophysiological experiments and *post hoc* imaging to confirm M4 ipRGC cell identity.

**Figure S12. Brn3b tunes the excitability of M4 ipRGCs.** (**A-B**) Individual traces of evoked firing response to current injections in control (A) and Brn3bcKO (B) M4 ipRGCs (n = 5-7 cells/group). (**C-D**) Individual traces of evoked firing response at different current steps (C) and current density (D) recorded in control and Brn3bcKO M4 ipRGCs. (**E**) Difference of voltage between baseline and after the current injection step control and Brn3bcKO in M4 ipRGCs. Data are Mean ± SEM, *P<0.05, two-way ANOVA with repeated measures.

**Figure S13. Brn3b does not affect the intrinsic photocurrent of M4 ipRGCs.** (**A**) Individual traces of photocurrent responses in control (top), Brn3bcKO (middle) and average photocurrent (bottom) of M4 ipRGCs (n = 4-7 cells/group). (**B**) Average current density of control and Brn3bcKO M4 ipRGCs. (**C**) Intrinsic photocurrent quantification of control and Brn3b M4 ipRGCs at different phases of the response. The left panel shows the Early, Intermediate and Late components of the photoresponses. Data are Mean ± SEM, n.s. (not significant, P>0.05, Mann Whitney U test.

**Figure S14. Brn3bcKO mice showed decreased contrast sensitivity.** (**A**) contrast sensitivity quantification in OKR at 0.031, 0.064, 0092 and 0.272 cycles/degree (c/d) in control and Brn3bcKO mice (n = 6-12 / group). (**B**) individual traces of contrast sensitivity vs. spatial frequencies in control and Brn3bcKO mice. Data are Mean ± SEM, n.s. (not significant) P>0.05, *P<0.05, **P<0.01, two-way ANOVA with repeated measures.

**Figure S15. Time course of pupillary light response (PLR) in control and Brn3bcKO mice.** (**A** and **C**) Pupil constriction plotted as a function of time in control (black) and Brn3bcKO (green) mice in response to 5 second dim (13.8 log quanta/cm2/s, A) and bright (14.8 log quanta/cm2/s, C) light stimuli. (**B** and **D**) Grouped data of the time constant Tau measured by fitting individual PLR data using a single-exponential decay function. There were no significant differences in Tau between control and Brn3bKO animals in response to dim (B) and bright (D) light stimuli. n.s. (not significant) P>0.05, Mann-Whitney U test.

**Figure S16. Period and activity profile parameters in control and Brn3bcKO.** (**A-B**) Brn3bcKO did not show different period length (A) or overall activity changes (B) compared to control mice (n = 6 / group). (**C-D**) The average activity onset was earlier (C) and more variable (D) in Brn3bcKO mice compared to control. (**E**) The activity onsets during 6-hour phase advance were not different in Brn3bcKO mice. Data are Mean ± SEM, n.s. (not significant) P>0.05, *P<0,05, **P<0.01, Student’s t and Mann Whitney U tests.

**Figure S17. Representative actograms from control and Brn3bcKO mice.** Representative wheel running activity profiles from control (top) and Brn3bcKO (bottom) mice.

**Figure S18. Characterization of *Opn4^Cre/+^ ; Rpl22^HA^* and *Opn4^Cre/+^ ; Brn3b^cKOAP^ Rpl22^HA^* mouse lines.** (**A**) (Top) HA expression in the ganglion cell layer (GCL) in control and Brn3bcKO mice. (Bottom) Brn3bcKO mice presented significantly lower number of HA-expressing RGC compared to control littermates (n = 5-6 /group). (**B**) (Top) HA expression in the outer nuclear layer (ONL) in control and Brn3bcKO mice (n = 5-6 /group). (Bottom) control and Brn3bcKO mice presented non-significant differences in the density of cone photoreceptors expressing HA. All data are Mean ± SEM, n.s. (not significant) P>0.05, **P<0.01, Mann Whitney U test.

**Figure S19. Pattern labeling of Brn3b and Opn4 mRNA in cross-sectioned retinas.** Brn3b mRNA probe (left, cyan) is present in the ganglion cell layer (GCL) in ipRGCs (full ellipse), other RGCs (dashed line ellipse) and sparse puncta in the inner plexiform layer (IPL). Opn4 mRNA probe (Right, magenta) is present only in the GCL in ipRGCs (full ellipse). INL: inner nuclear layer; OPL: outer plexiform layer; ONL: outer nuclear layer; OS: outer segments of the cone and rod photoreceptors; RPE: retinal pigmented epithelium.

